## Supplemental figures for "Disordered regions and folded modules in CAF-1 promote histone deposition in *S. pombe*"

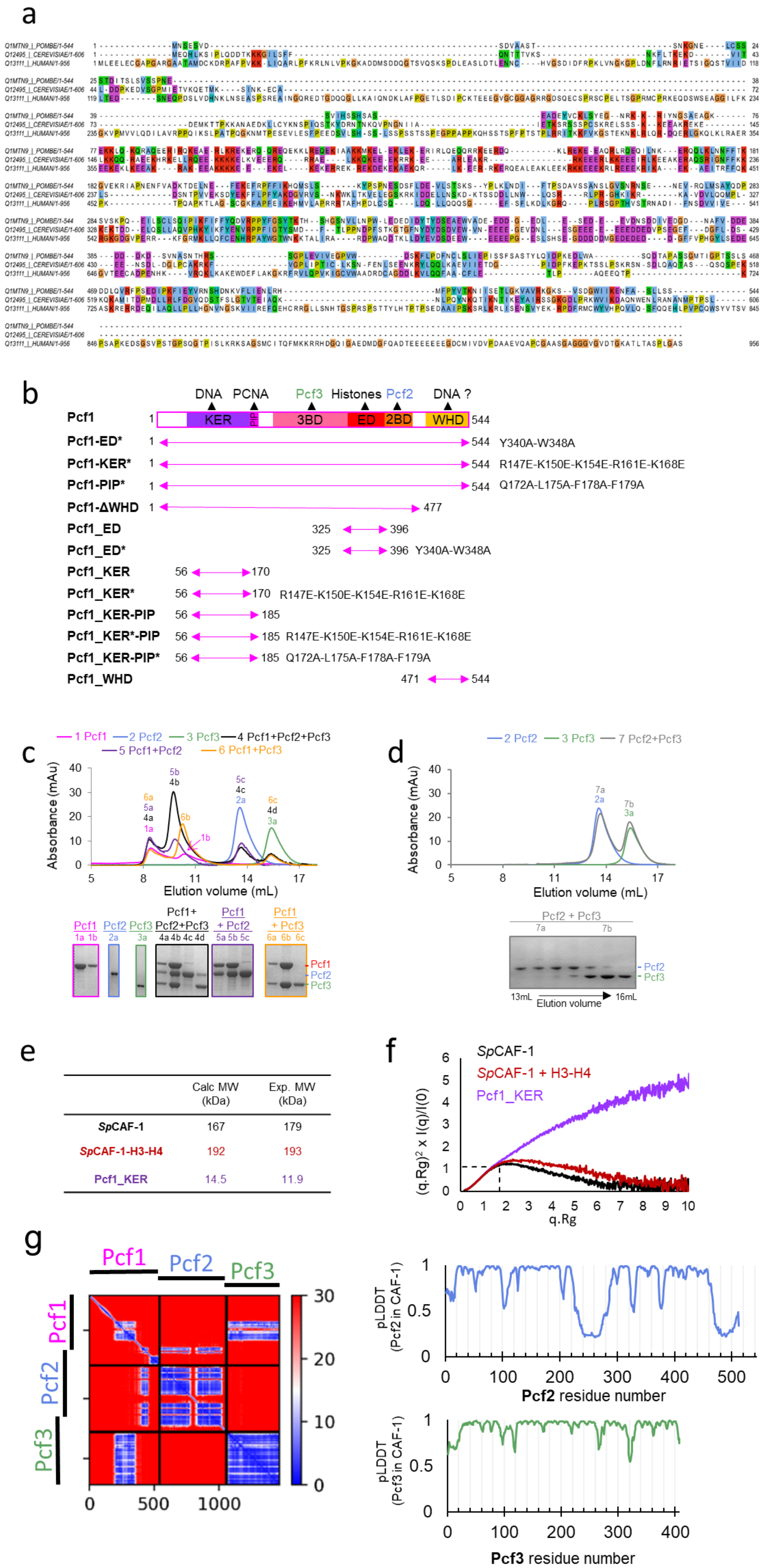
a
Q1MTN9_|_POMBE/1-544
Q12495_|_CEREVISIAE/1-606
Q13111_|_HUMAN/1-956
b
DNA PCNA Pcf3 Histones Pcf2 DNA ?
Pcf1 1 KER PIP 3BD ED 2BD WHD 544
Pcf1-ED* 1 544 Y340A-W348A
Pcf1-KER* 1 544 R147E-K150E-K154E-R161E-K168E
Pcf1-PIP* 1 544 Q172A-L175A-F178A-F179A
Pcf1-ΔWHD 1 477
Pcf1_ED 325 396
Pcf1_ED* 325 396 Y340A-W348A
Pcf1_KER 56 170
Pcf1_KER* 56 170 R147E-K150E-K154E-R161E-K168E
Pcf1_KER-PIP 56 185
Pcf1_KER*-PIP 56 185 R147E-K150E-K154E-R161E-K168E
Pcf1_KER-PIP* 56 185 Q172A-L175A-F178A-F179A
Pcf1_WHD 471 544
c
1 Pcf1 2 Pcf2 3 Pcf3 4 Pcf1+Pcf2+Pcf3
5 Pcf1+Pcf2 6 Pcf1+Pcf3
Absorbance (mAu)
Elution volume (mL)
Pcf1 Pcf2 Pcf3 Pcf1+ Pcf2+Pcf3 Pcf1 + Pcf2 Pcf1 + Pcf3
Pcf1 Pcf2 Pcf3
d
2 Pcf2 3 Pcf3 7 Pcf2+Pcf3
Absorbance (mAu)
Elution volume (mL)
Pcf2 + Pcf3
13mL Elution volume 16mL
e
| | Calc MW (kDa) | Exp. MW (kDa) |
| SpCAF-1 | 167 | 179 |
| SpCAF-1-H3-H4 | 192 | 193 |
| Pcf1_KER | 14.5 | 11.9 |
f
SpCAF-1
SpCAF-1 + H3-H4
Pcf1_KER
(q.Rg)² x I(q)/I(0)
q.Rg
g
Pcf1 Pcf2 Pcf3
pLDDT (Pcf2 in CAF-1)
Pcf2 residue number
pLDDT (Pcf3 in CAF-1)
Pcf3 residue number

**Figure S1: General organization of the full *Sp*CAF-1 complex. a** Alignment of the large-subunit of CAF-1 from *S. pombe*, *S. cerevisiae* and *human* **b** Domain composition of Pcf1 and binding predictions inferred from sequence homology. The delimitation and mutant positions of the different constructs of Pcf1 produced for this study are shown. **c-d** SEC analysis of recombinant Pcf1, Pcf2 and Pcf3 proteins purified separately (1-3) mixed by pairs (5-7) or the mix of the three proteins (4). For each chromatograph, the composition of all peaks (identified with the run number (1-7) and the peak position (a-d)) were analysed by SDS-PAGE. **e** Molecular weight calculated with SAXS data (Exp. MW) compared to theoretical MW (Calc. MW) for *Sp*CAF-1 Pcf1:Pcf2:Pcf3 1:1:1, for *Sp*CAF-1-H3-H4 Pcf1:Pcf2:Pcf3:H3:H4 1:1:1:1:1 and monomeric Pcf1_KER. **f** Dimensionless Kratky plot of the SAXS experiment of *Sp*CAF-1 (black) and *Sp*CAF-1−H3−H4 (red) and Pcf1_KER (purple). The position of the expected maximum of the curve for a fully globular protein is shown with dashed black lines. The curve of Pcf1_KER does not tend towards zero for large q.Rg values, showing that this domain has an extended structure. The position of the maximum for the curve obtained for *Sp*CAF-1−H3−H4 is shifted to higher values in the y and x axis compared to the *Sp*CAF-1 curve. This indicates that the *Sp*CAF-1−H3−H4 complex is more extended that *Sp*CAF-1 alone. Further interpretations are given by the fit of experimental intensities with models from AlphaFold (see **Figure S1m-p**). **g** Left panel: Predicted Alignment Error plot (PAE) calculated by AlphaFold2 for the best model for the full *Sp*CAF-1 complex. Right panels: Local Distance Difference Test (pLDDT) calculated by the AlphaFold2 for Pcf2 (upper right) and Pcf3 (lower right) for this model. pLDDT for Pcf1 is shown in **Figure 1b**.

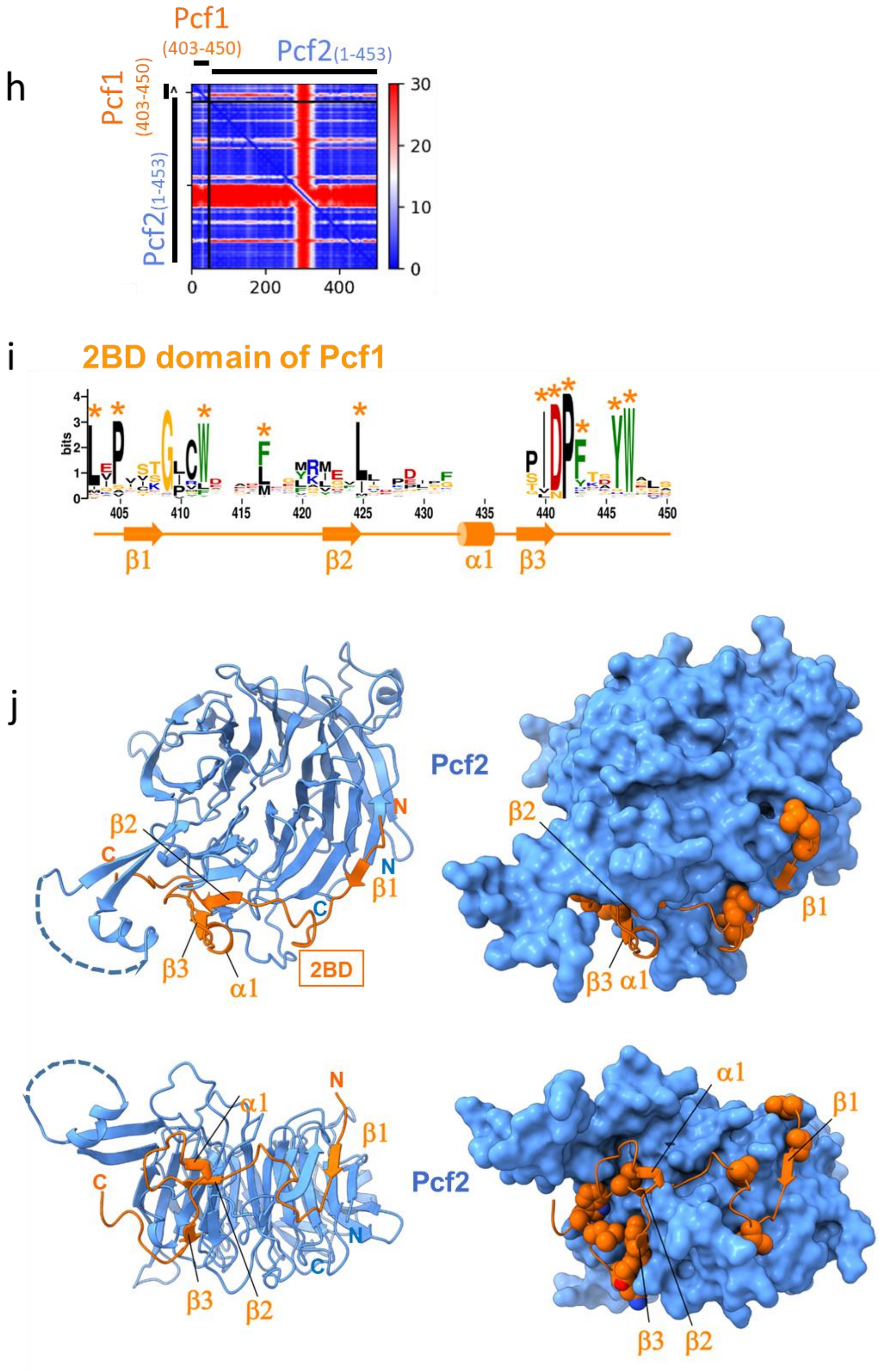


**Figure S1 (continued) h** Predicted Alignment Error plot (PAE) calculated by AlphaFold2 for the best model for the Pcf1(403-450)-Pcf2(1-453) complex corresponding the module of *Sp*CAF-1 including the seven WD repeats of Pcf2 and the segment of Pcf1 covering the 2BD domain **i** Sequence logo of the 2BD region of Pcf1 (Pcf1_2BD). Residues indicated with a star are correspond to conserved residues at the interface, they are shown as spheres in h. The secondary structure in the model is shown below the residue numbering. **j** Best AlphaFold2 model for this module with two perpendicular orientations (upper panels and lower panels respectively). Pcf2 is shown in blue and Pcf1_2BD in orange. In the left panels, the module is presented with cartoons and in the right panels, Pcf2 is shown with a surface and Pcf1_2BD a cartoon with conserved residues highlighted with a star in **i** represented as spheres. A disordered loop (234-283) of Pcf2 is omitted and represented with a dashed line. The model is available https://www.modelarchive.org/doi/10.5452/ma-1bb5w.

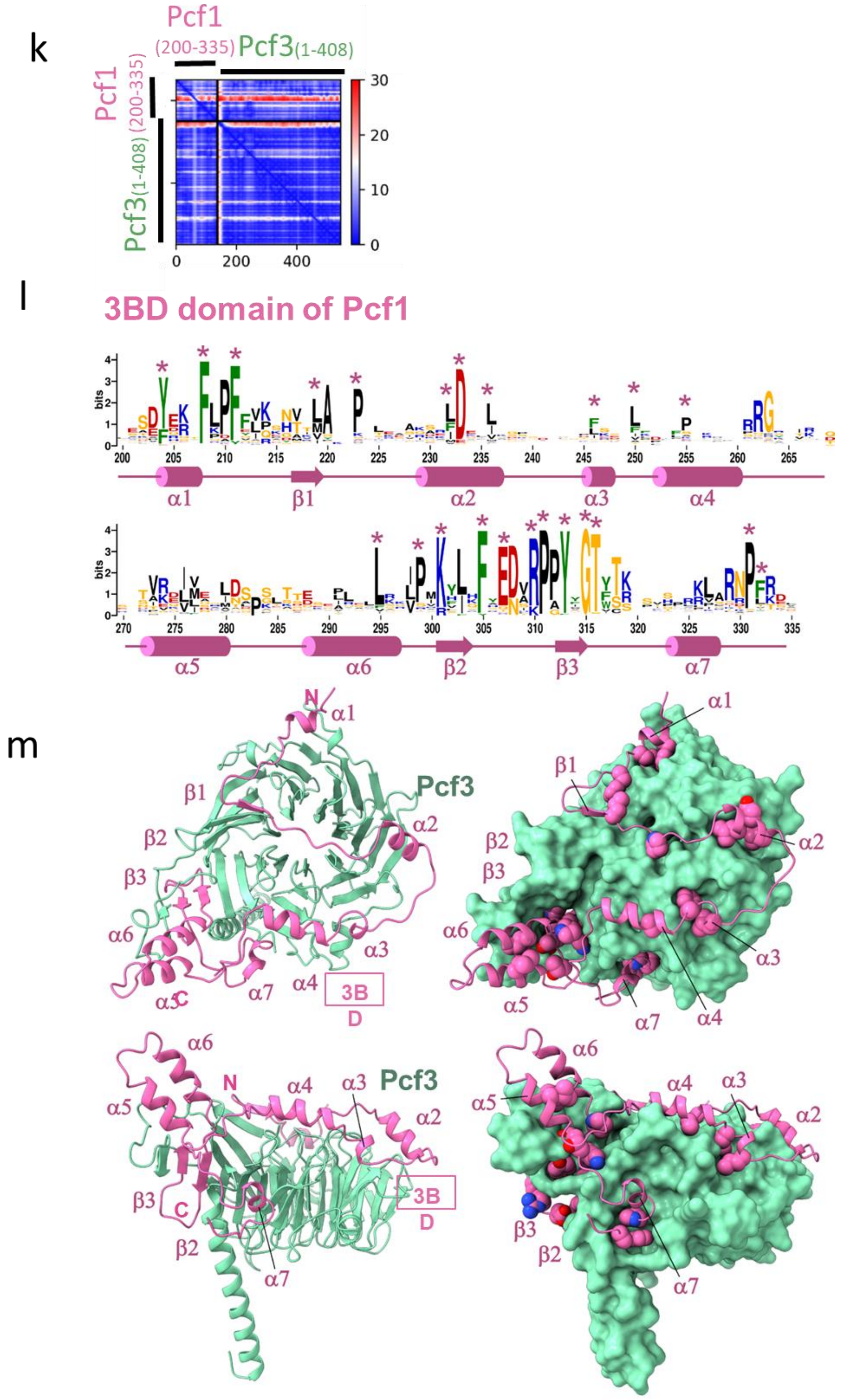


**Figure S1 (continued) : k** Predicted Alignment Error plot (PAE) calculated by AlphaFold2 for the best model for the Pcf1(200-335)-Pcf3(1-408) complex corresponding the module of *Sp*CAF-1 including the seven WD repeats of Pcf3 and the segment of Pcf1 covering the 3BD domain. **l** Sequence logo of the 3BD region of Pcf1 (Pcf1_3BD). Residues indicated with a star are correspond to conserved residues at the interface, they are shown as spheres in l. The secondary structure in the model is shown below the residue numbering. **m** Best AlphaFold2 model of this module corresponding to the subunit Pcf3 and Pcf1_3BD shown in **l** with two perpendicular orientations (upper panels an lower panels respectively). Pcf3 is shown in green and Pcf1_3BD in pink. In the left panels, the module is presented with cartoons and in the right panels, Pcf3 is shown with a surface and Pcf1_3BD a cartoon with conserved residues highlighted with a star in **l** represented as spheres. The model is available https://www.modelarchive.org/doi/10.5452/ma-bxxkp

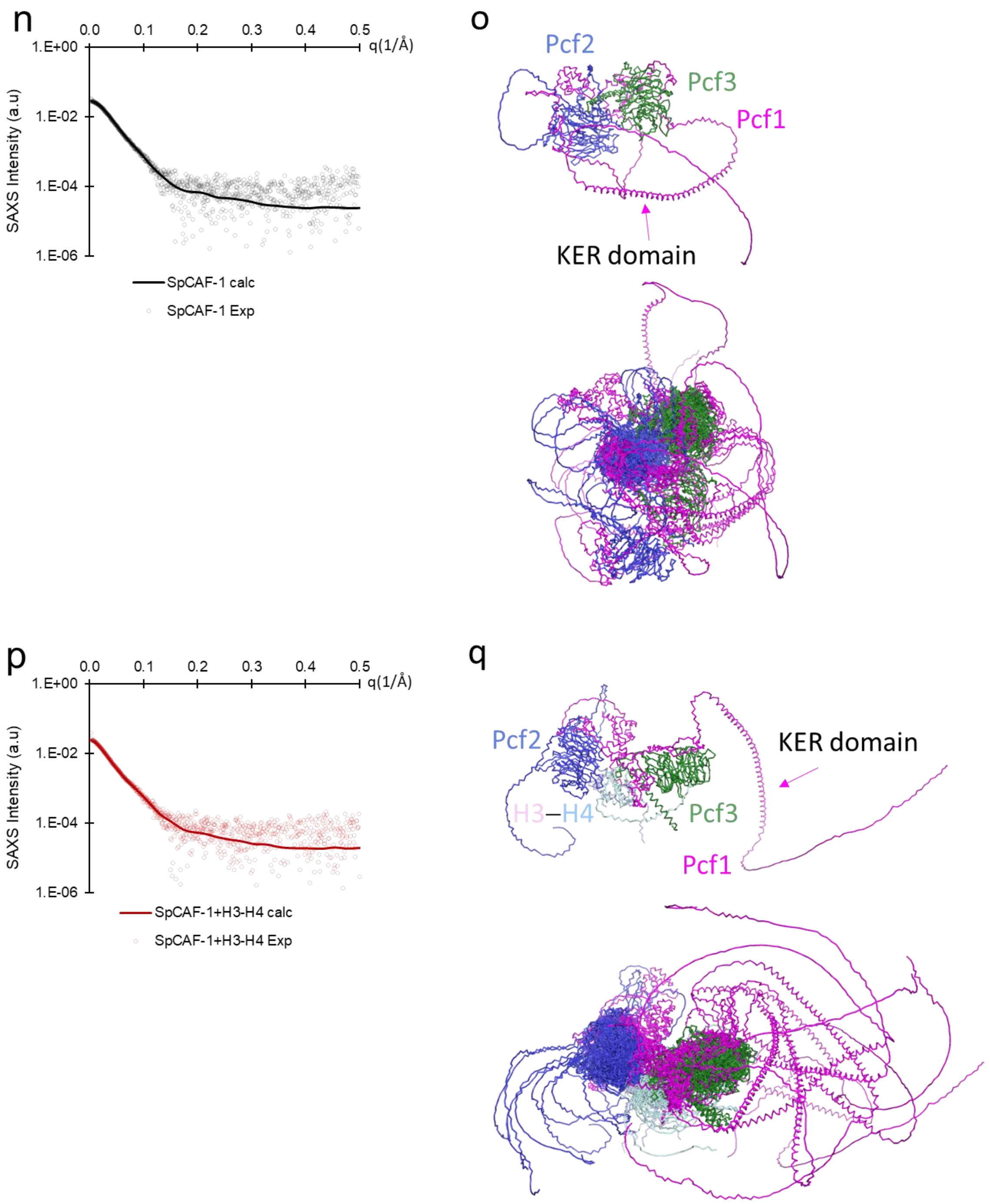


**Figure S1 (continued) : n-p** Experimental and fitted SAXS profile intensity (I) as a function of the momentum transfer (q) for *Sp*CAF-1 (black circles) and for the best model generated by the Dadimodo Software(Rudenko et al. 2019) (continuous black line) shown in **o** (upper panel) . **o** Ribbon representation of models generated by the Dadimodo software(Rudenko et al. 2019) fitting data in **n**. Upper panel: Best model. Lower panel: overlay of the 8 best models. **o** Experimental and fitted SAXS profile intensity (I) as a function of the momentum transfer (q) for *Sp*CAF-1−H3−H4 (red circles) and for the best model generated by Dadimodo Software (continuous red line) shown in **q** (upper panel). **q** Ribbon representation of models generated by the Dadimodo software fitting data in **o**. Upper panel: Best model. Lower panel: overlay of the 8 best models. **o-q**, Pcf1 is shown in magenta, Pcf2 in blue and Pcf3 in green and H3−H4 in light pink and turquoise respectively. SAXS experimental and fitting details are given in the Methods section and **Table S1**.

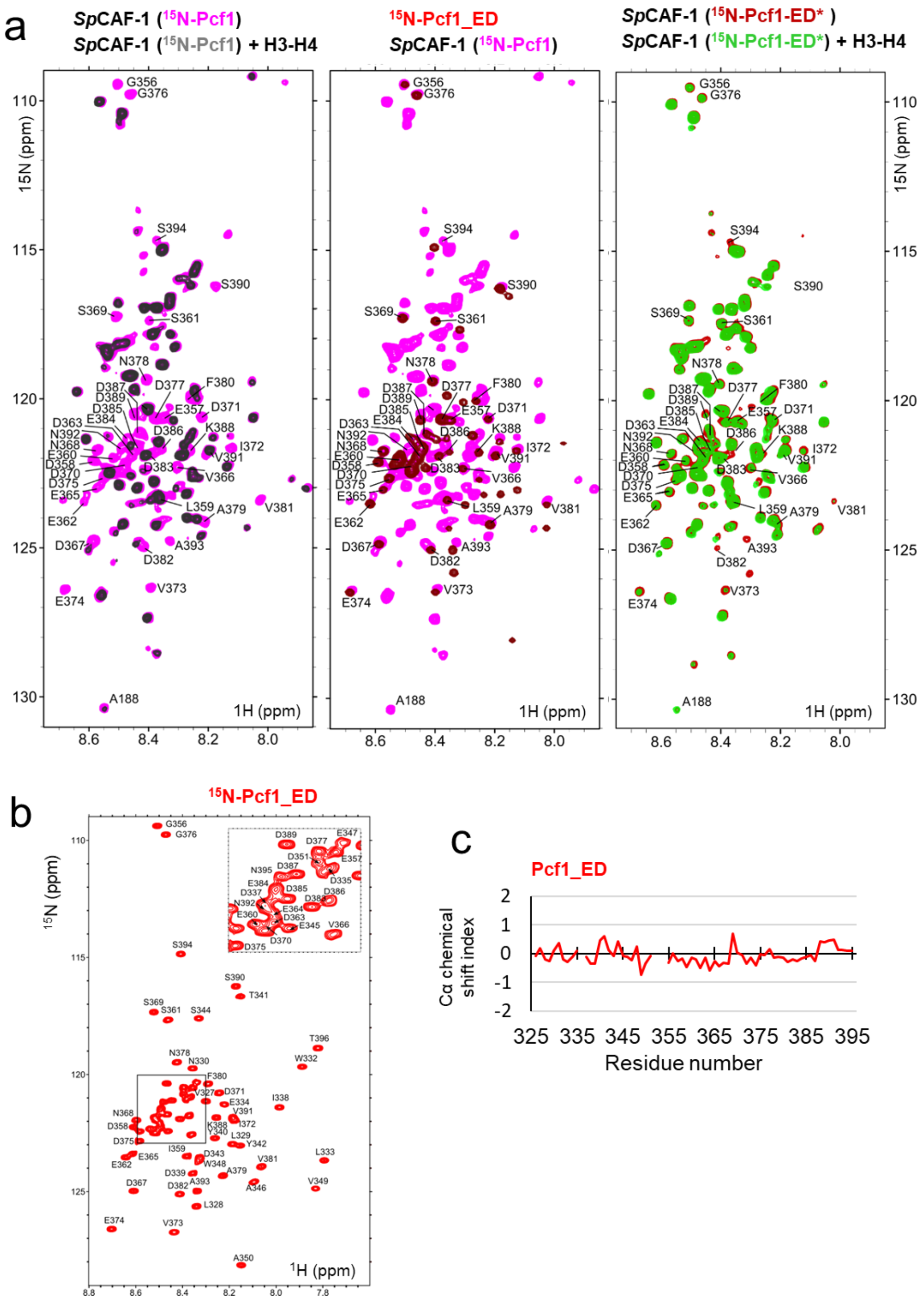


**Figure S2: The acidic ED domain binds histones alone and in the full CAF-1 complex. a** left panel : Overlay of the NMR $^{1}$H-$^{15}$N SOFAST-HMQC spectrum of *S. pombe* CAF-1($^{15}$N-Pcf1) alone (magenta) and after addition of *S. pombe* H3−H4 (1:1) (black). middle panel: Overlay of the $^{1}$H-$^{15}$N SOFAST-HMQC spectrum of *S. pombe* CAF-1($^{15}$N-Pcf1) alone (magenta) with the $^{1}$H-$^{15}$N Pcf1_ED spectrum in the same buffer conditions (red). right panel : Overlay of the $^{1}$H-$^{15}$N SOFAST-HMQC spectrum of *S. pombe* CAF-1($^{15}$N-Pcf1-ED*) alone (red) and after addition of *S. pombe* H3−H4 (1:1) (green). The assignments of residues from IDR3 are indicated in all panels. **b** $^{1}$H-$^{15}$N HSQC spectra of Pcf1_ED (325-396). The assignments of the central area of the spectrum are zoomed in the box at the top right of the spectrum. Spectral dispersion of HN resonances, ranging from 8.1 to 8.6 ppm are consistent with all residues being mainly disordered **c** Cα chemical shift index of the isolated Pcf1-ED domain. Values close to zero of the Cα chemical shift index confirms the unfolded nature of the ED_domain. All spectra were recorded at 10°C.

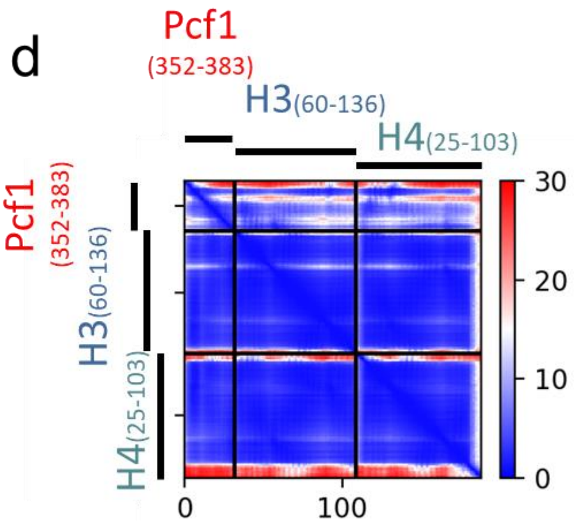


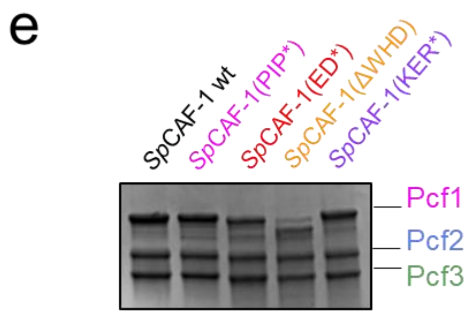


**Figure S2 (continued) : d** Predicted Alignment Error plot (PAE) calculated by AlphaFold2 for the best model for the Pcf1(352-383)-H3(60-136)-H4(25-103). **e.** SDS-PAGE electrophoresis of the reconstituted CAF-1 complexes WT and mutants used in this study. The model is available https://www.modelarchive.org/doi/10.5452/ma-htx0n.

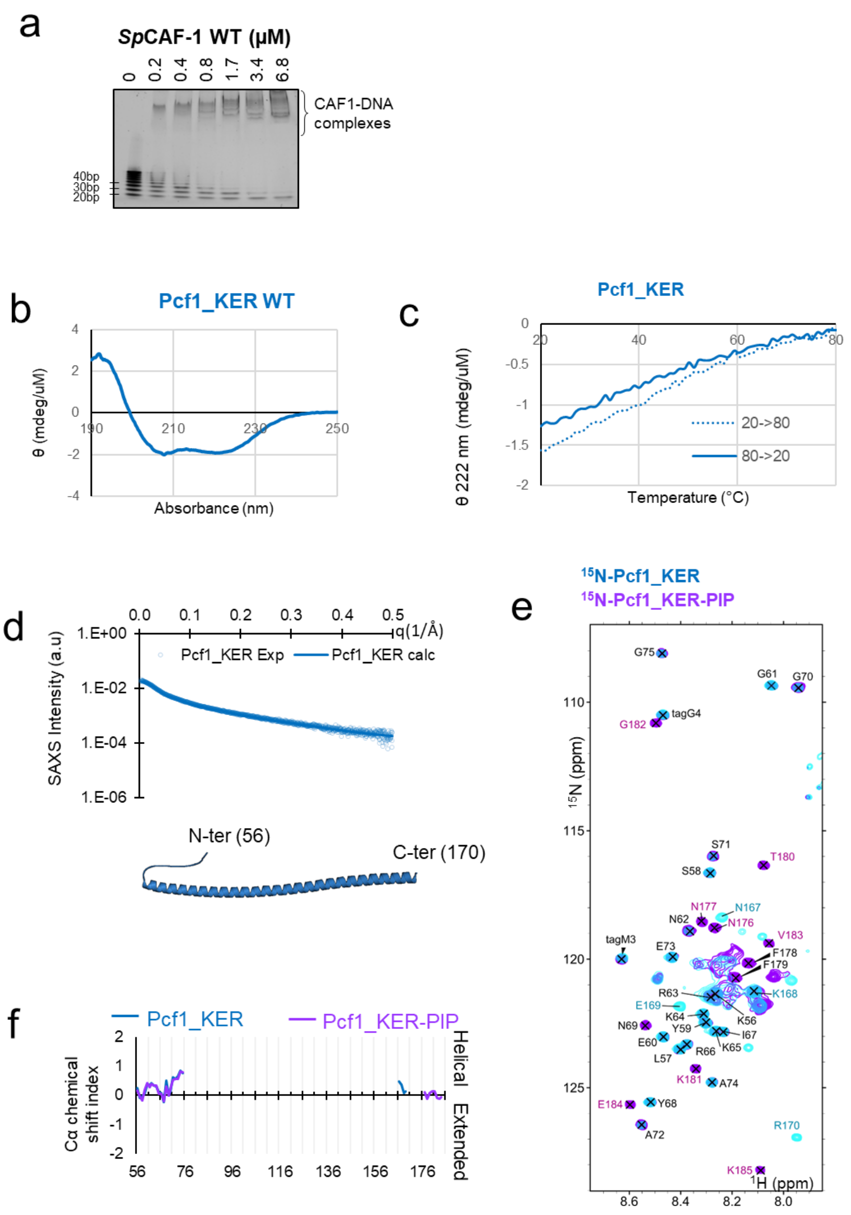


**Figure S3: Pcf1_KER is the main DNA binding domain of *S. pombe* CAF-1. a** EMSA with *Sp*CAF-1 and a ladder of 20–100 bp dsDNA fragments revealed with SYBR SAFE staining. **b** Circular dichroism (CD) spectrum of the Pcf1_KER domain (56-170) at 20°C. The measured signal confirms the high helical content of this fragment. **c** Evolution of the CD measurements of Pcf1_KER at 222nm during a ramp temperature from 20°C to 80°C (forth with a dashed line and back with a continuous line). The linear unfolding without any cooperativity is consistent with the absence of tertiary structure of this domain. **d** Upper panel: Experimental SAXS profile intensity (I) as a function of the momentum transfer (q) for Pcf1_KER (cyan circles) and fitted (continuous blue line). Lower panel : Cartoon representation of the best model fitting the experimental SAXS data. Fitting details are given in Methods section and **Table S1**. The models are in very good accuracy with the AF2 models of a strait helix. **e** Overlay of the $^{1}$H-$^{15}$N SOFAST-HMQC spectrum of $^{15}$N-Pcf1_KER (cyan) and $^{15}$N-Pcf1_KER-PIP (purple) at 10°C. Assigned resonances are indicated in black for signals that overlap for both constructs, in blue for resonances observed only for $^{15}$N-Pcf1_KER and in purple for residues observed only for $^{15}$N-Pcf1_KER-PIP. For $^{15}$N-Pcf1_KER (cyan), as expected for a long anisotropic helix , only the few signals that were assigned to its 20 first and 4 last residues were visible, , whereas helical residues are not visible in the spectrum. For the $^{15}$N-Pcf1_KER-PIP (purple), we did not observe the NMR signals for all residues PIP box, but only starting at $N_{176}$, suggesting that, the helix could extends towards the N-ter part of the PIP motif. **f** Cα chemical shift index of the assigned residues for Pcf1_KER (cyan) and Pcf1_KER-PIP (purple) are consistent with disordered ends of the segments, with some helical propensity at the N-and C-terminus of the long helix. Together these results are in perfect adequation with models predicted by AF2.

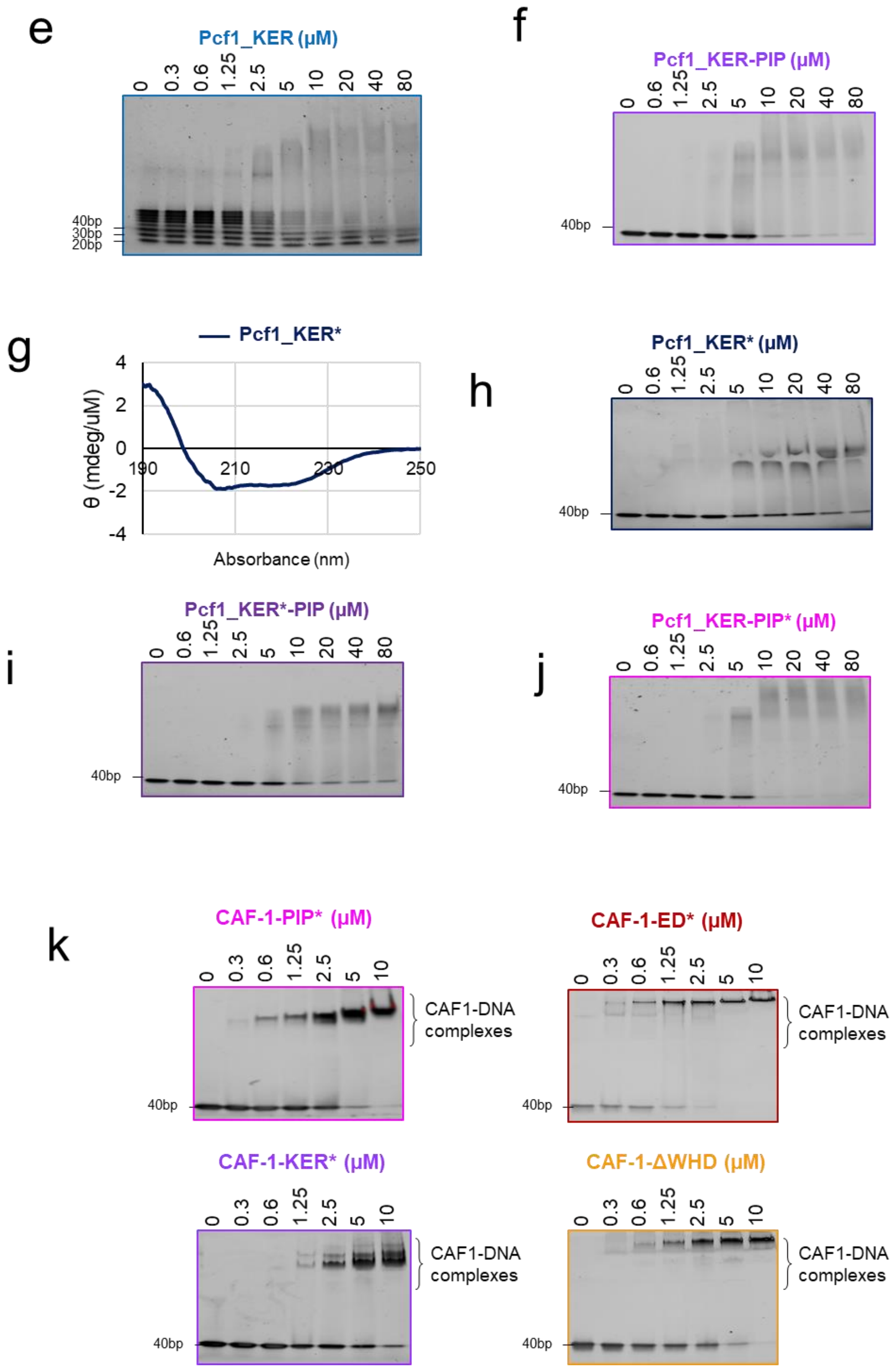


**Figure S3 (continued): e** EMSA showing Pcf1_KER binding to a 20–100 bp ladder of ds DNA fragments. **f** EMSA showing Pcf1_KER-PIP binding to a 40bp dsDNA. **g** CD spectra of the mutated Pcf1_KER* at 20°C. **h-j** EMSA showing the binding with a 40bp dsDNA of Pcf1_KER* (h), Pcf1_KER*-PIP (i) and Pcf1_KER-PIP* (j). **k** EMSA showing the full *Sp*CAF-1 binding to a 40 dsDNA revealed with SYBR SAFE staining.

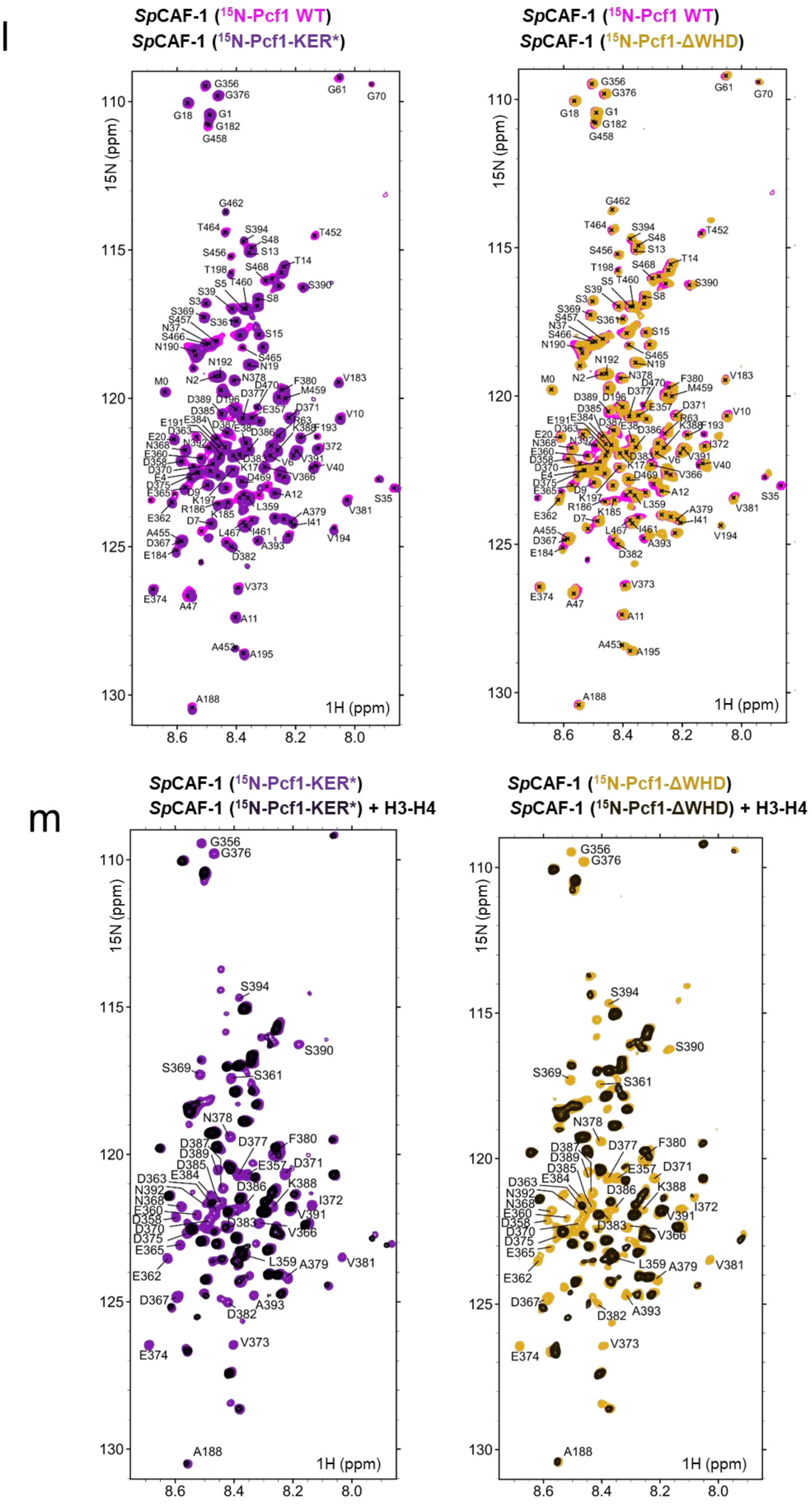


**Figure S3 (continued): l** Left panel: Overlay of the $^{1}$H-$^{15}$N SOFAST-HMQC spectrum of *Sp*CAF-1($^{15}$N-Pcf1) alone (magenta) and *Sp*CAF-1($^{15}$N-Pcf1-KER*) (purple). Right panel: Overlay of the $^{1}$H-$^{15}$N SOFAST-HMQC spectrum of *Sp*CAF-1($^{15}$N-Pcf1) alone (magenta) and *Sp*CAF-1($^{15}$N-Pcf1-ΔWHD) (orange). The assignments of residues in *Sp*CAF-1($^{15}$N-Pcf1) are indicated. **m** Left panel: Overlay of the $^{1}$H-$^{15}$N SOFAST-HMQC spectrum of *Sp*CAF-1($^{15}$N-Pcf1-KER*) alone (purple) and after addition of *Sp*H3−H4 (1:1) (black). Right panel: Overlay of the $^{1}$H-$^{15}$N SOFAST-HMQC spectrum of *Sp*CAF-1($^{15}$N-Pcf1-ΔWHD) alone (orange) and after addition of *Sp*H3−H4 (1:1) (black).

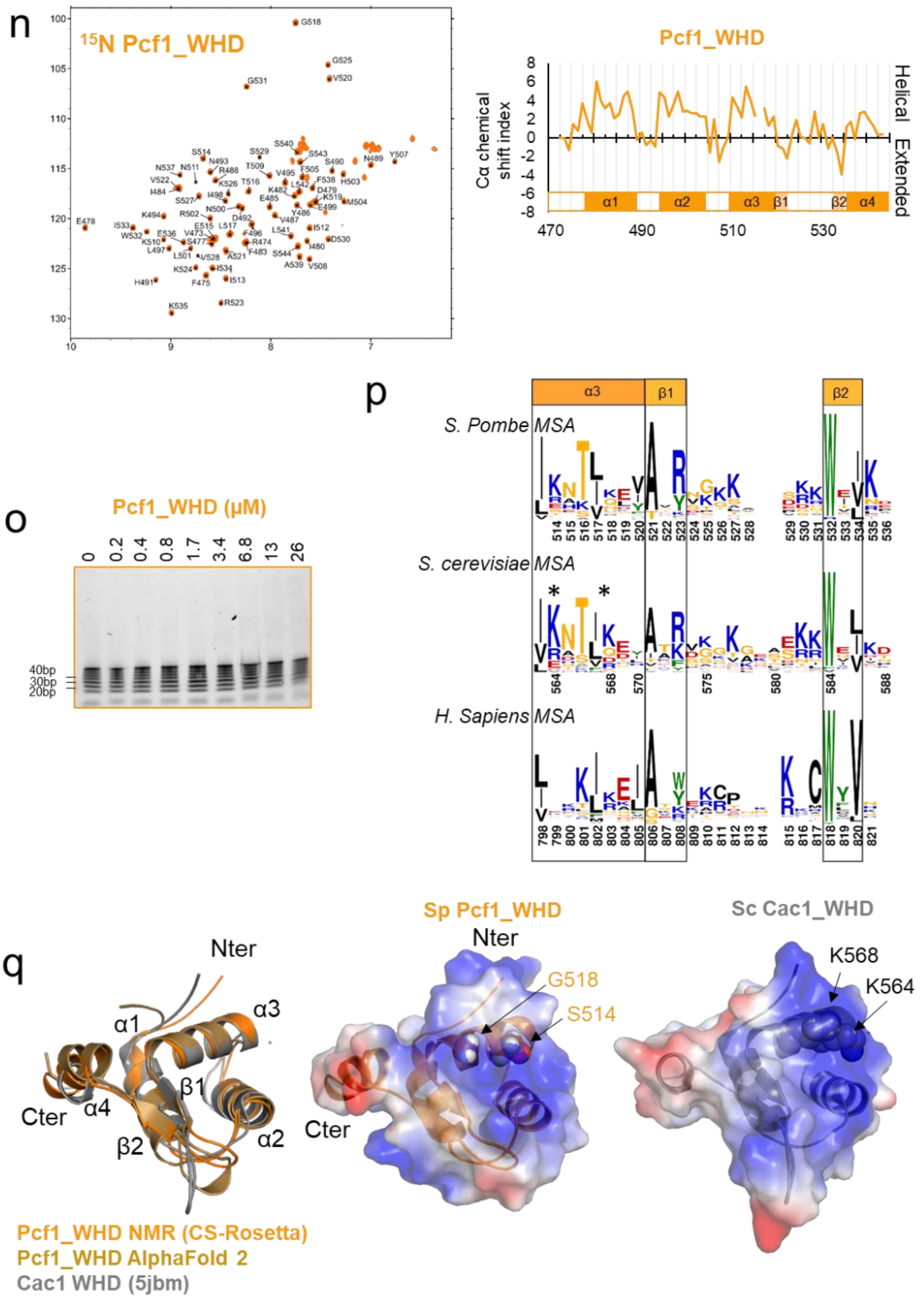


**Figure S3 (continued): Pcf1_WHD domain adopts a WHD fold similar to Sc C-terminal domain of Cac1, but does not bind DNA alone. n** Left panel: $^{1}$H-$^{15}$N HSQC spectrum of the Pcf1_WHD domain at 10°C. Assignment of are indicated. The large spectral dispersion HN signal is consistent with a fully folded domain Right panel : Cα chemical shift index of Pcf1_WHD. Positive values are observed in the four helices and negative values in two short beta strands. These secondary structures are consistent with the AF2 model. **o** EMSA revealed with SYBR SAFE staining with Pcf1_WHD domain and a ladder of 20–100 bp dsDNA. **p** Sequence Logo covering helix α3 and the two beta strands of WHD generated with three sequence datasets closed to *S. pombe* Pcf1, *S. cerevisiae* Cac1 and *H sapiens* CHAF1A/p150, respectively. Alignment of sequences was done based on structure superimposition. The two residues involved in DNA binding of Sc Cac1_WHD are highlighted with stars. **q** Left panel: Overlay of the calculated model of the WHD domain with CS-rosetta software (light orange), with AlphaFold2 (gold) and the structure of Cac1 WHD from budding yeast (PDB 5jbm, in grey) (Grey). Middle and right panels: *Sp*Pcf1_WHD and *Sc*Cac1_WHD, respectively, with their electrostatic potential (same orientation as in the left panel). The two basic residues responsible for DNA binding of *Sc*Cac1_WHD are highlighted with spheres and labeled, the corresponding residues in *Sp*Pcf1_WHD are also highlighted and labeled. Differences in the electrostatic surfaces may explain why Pcf1_WHD does not interact with DNA.

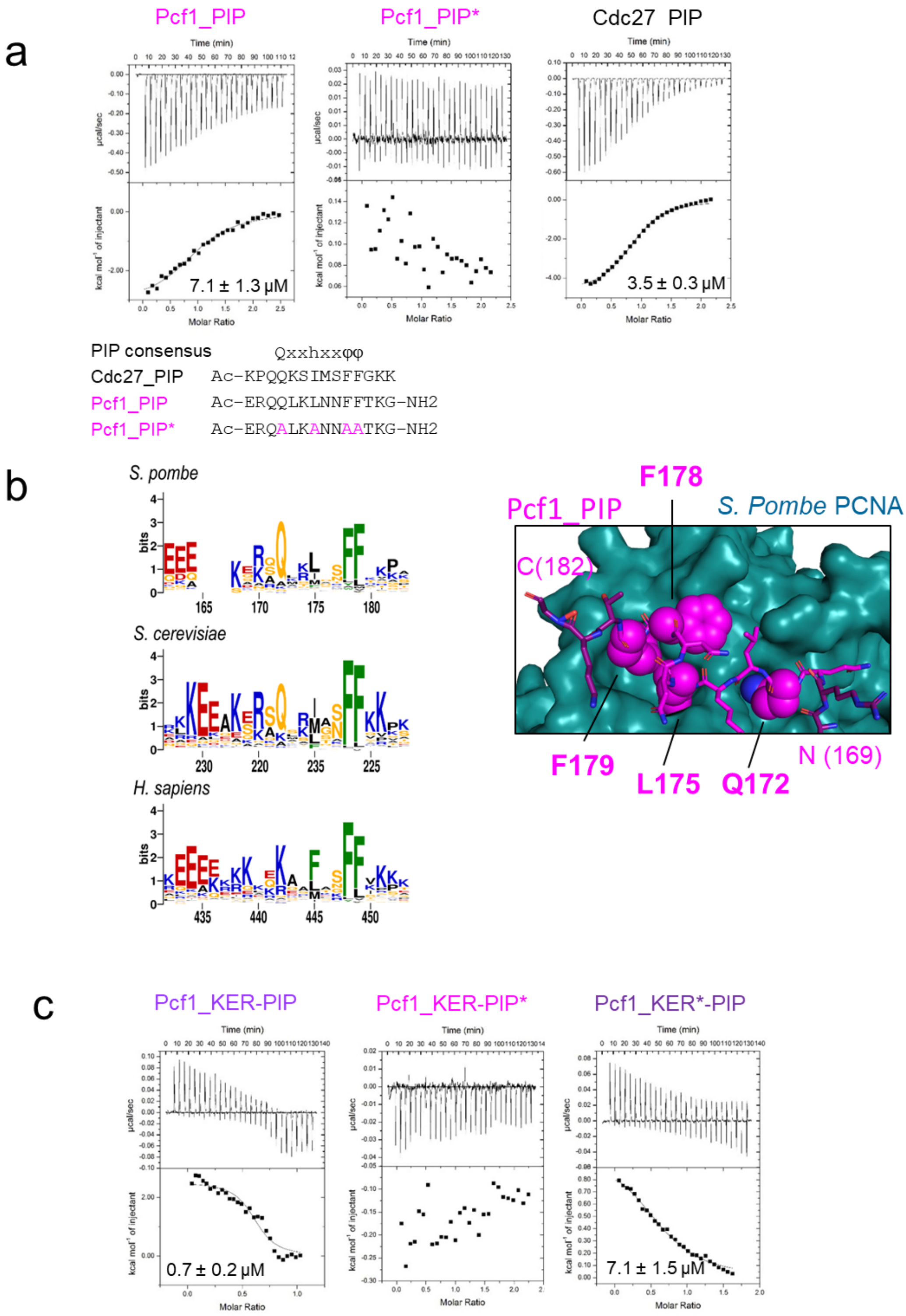


**Figure S4: The CAF-1 KER*mutant is affected for PCNA binding. a** ITC thermograms and data fitting for the indicated peptides upon titration of SpPCNA. The sequence of the three peptides is indicated as well as the canonical consensus for a PIP motif (h means hydrophobic residue, and φ an aromatic residue). **b** Left panel : Sequence Logo generated with a sequence data set adjacent to *S. pombe* Pcf1, *S. cerevisiae* Cac1 and H sapiens CHAF1A/p150 PIP motif, located at the C-terminus of the KER domain. Right panel : AlphaFold 2 model of Pcf1_PIP peptide (magenta) bound to SpPCNA (shown as a blue surface). Residues of the motif are highlighted with spheres and labeled. **c** ITC thermograms and data fitting for the indicated construct upon titration of SpPCNA.

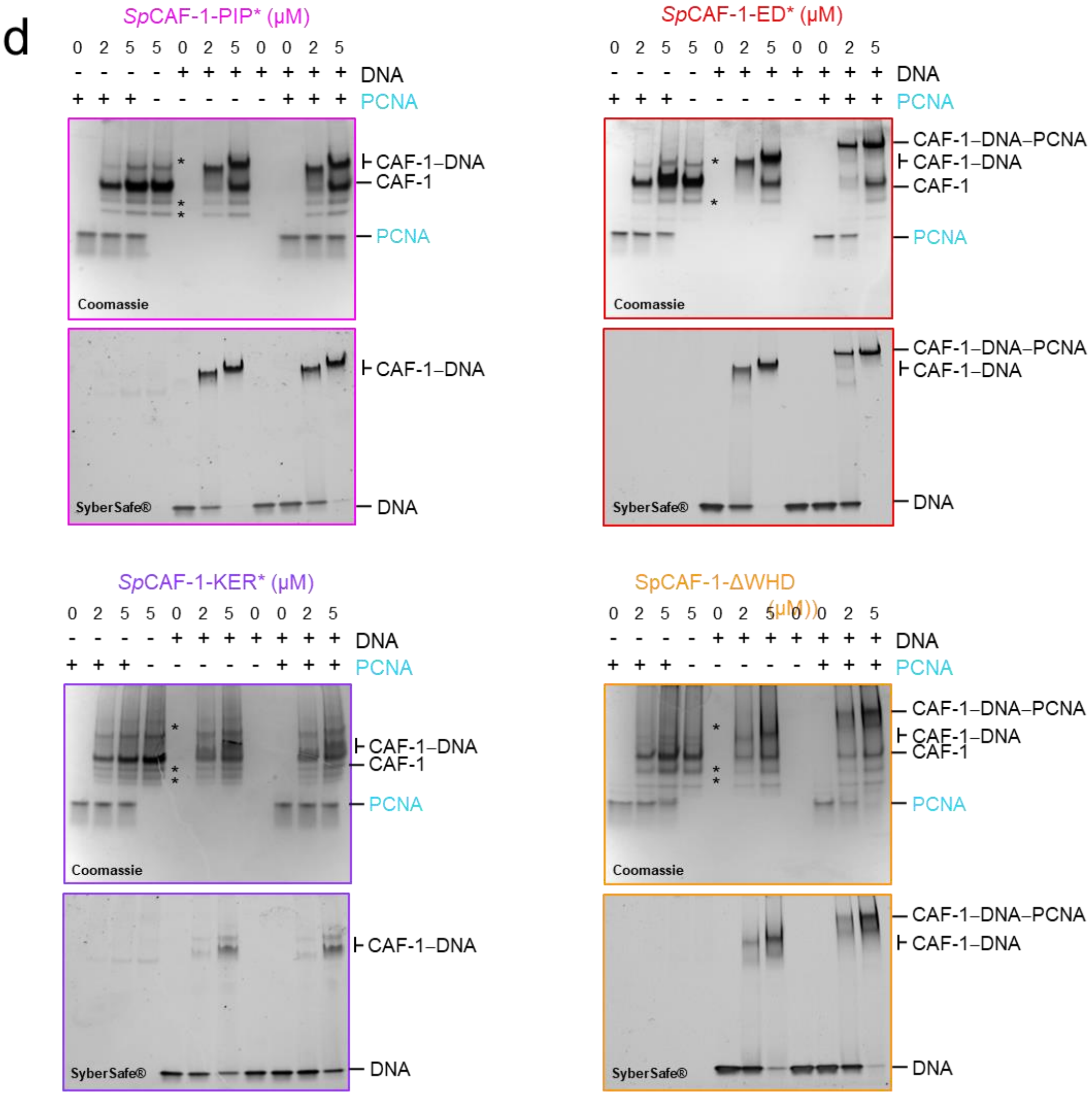


**Figure S4 (continued): The CAF-1 KER*mutant is affected for PCNA binding. d** EMSA showing interactions of purified CAF-1 mutated complexes (at the indicated concentrations), with or without recombinant SpPCNA (3µM) in the presence and absence of 40bp dsDNA (1µM). For each mutant, the revelation was done with Coomassie blue to reveal protein shifts in the upper panel, and with SYBR SAFE staining in lower panel to reveal DNA shifts.

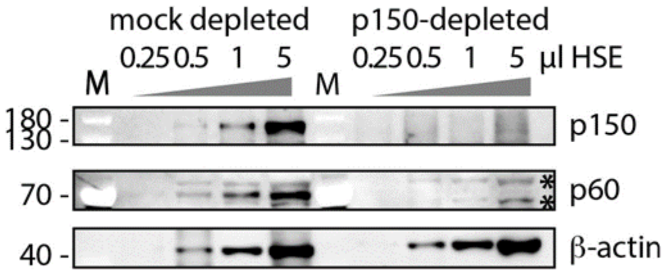


**Figure S5: Western blot analysis of mock- and p150-depleted HSE.** Xenopus p150 and p60 CAF-1 are shown for 0.25, 0.5, 1 and 3 μL of HSE as indicated. β-actin is used as a loading control. Stars on the p60 indicate non-specific bands. M, molecular weight markers indicated on the left**.**

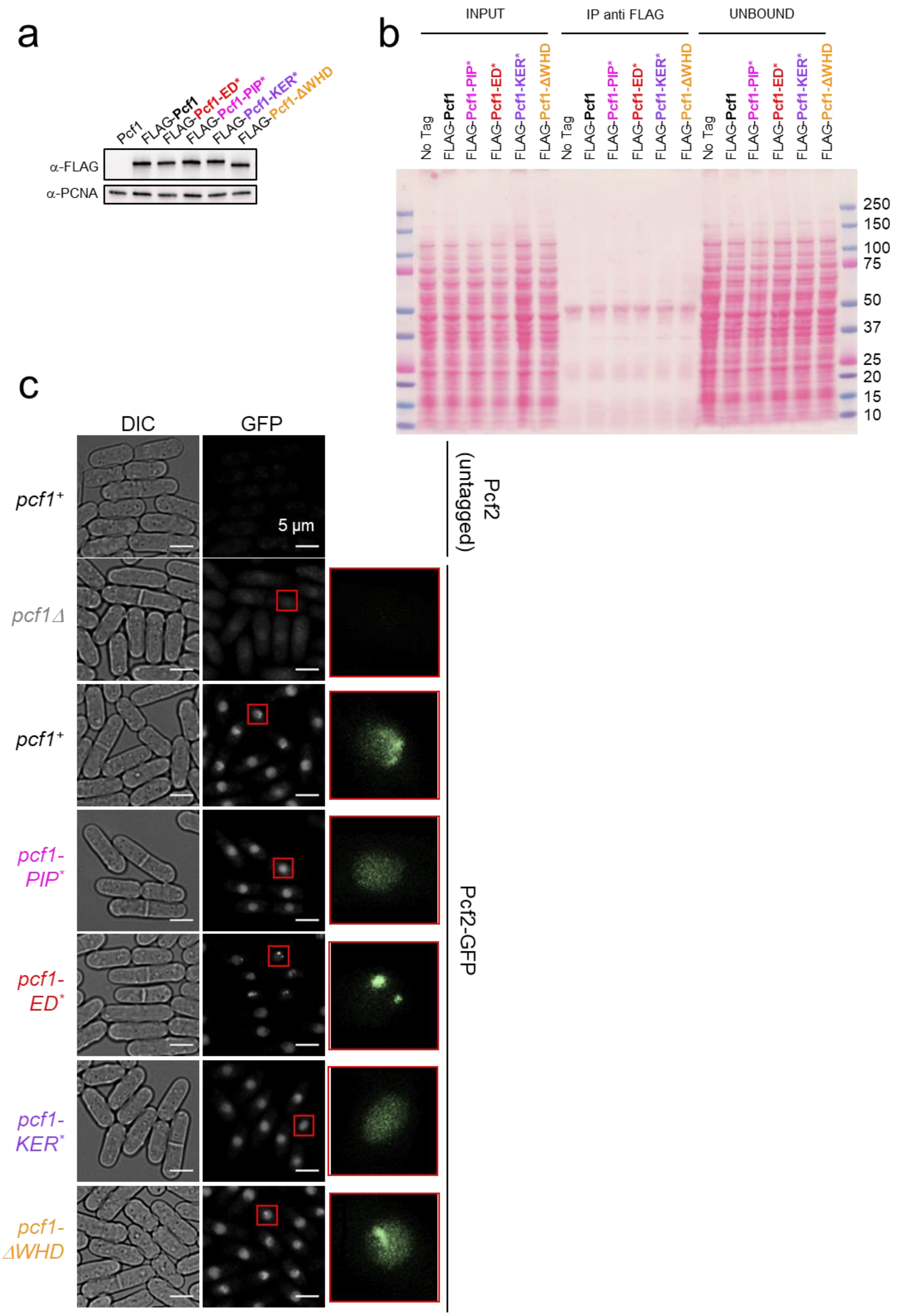


**Figure S6: Association of CAF-1 with histone is coupled to PCNA interaction *in vivo.*** **a** Expression levels of the FLAG-Pcf1 in indicated strains from total extracts. PCNA was used as loading control. **b** Nitrocellulose membrane from Figure 6A stained with Red ponceau. **c** Example of Pcf2-GFP foci in living cells in indicated strains. The top strain corresponds to a strain expressing wild-type and untagged Pcf2 as a negative control of GFP fluorescence. A zoom on a S-phase nuclei (from cells exhibited a septum) is showed for each strain.

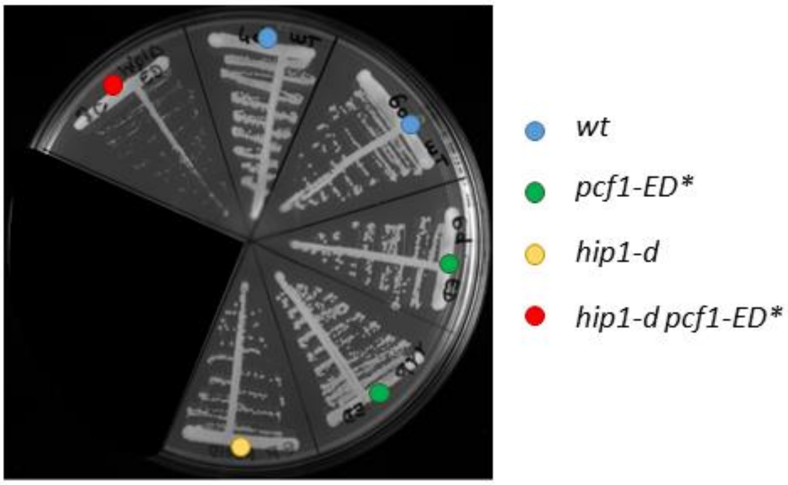


**Figure S7: the *pcf1-ED* mutation confers a synthetic growth defect when combined with hip1*Δ.** Spores of the indicated genotypes were streaked onto a YEA agar plate and grown at 30°C for 3 days.

### Table S1: Experimental information and modelling of SAXS data

| **Sample details** | **Pcf1_KER** | ***Sp*CAF-1** | ***Sp*CAF-1+H3-H4** |
|---|---|---|---|
| Organism | *S. Pombe* | | |
| Source (catalogue No. or reference) | Recombinant proteins (See Methods) | | |
| UniProt sequence ID (residues in construct) | Q1MTN9 | Q1MTN9-O13985- Q9Y825 | Q1MTN9-O13985-Q9Y825-P09988-P09322 |
| Extinction coefficient [$A_{280}$, 0.1%(*w/v*)] | 0.207 | 1.15 | 1.045 |
| M from chemical composition (Da) | 14 410 | 167 386 | 194 130 |
| SEC–SAXS column | S200 5/150 Increase | | |
| Loading concentration (mg $ml^{-1}$) | 0.7 | 3.3 | 2.6 |
| Injection volume (µl) | 45 | 50 | 50 |
| Flow rate (ml $min^{-1}$) | 0.3 | 0.3 | 0.3 |
| Solvent (solvent blanks taken from SEC flowthrough prior to elution of protein) | Tris 10 mM, NaCl 150 mM, β-mercaptoethanol 5 mM, pH 8 | | |
| **SAXS data-collection parameters.** | | | |
| Instrument/data processing | BioSAXS on the SWING beamline at Synchrotron SOLEIL(Thureau et al. 2021) | | |
| Wavelength (Å) | 1.0332 | | |
| Beam size (µm) | 500x200 | | |
| Camera length (m) | 2.00 | | |
| q measurement range ($Å^{-1}$); $q = 4\pi\sin(\theta)/\lambda$ (2θ: scattering angel & λ the x-ray wavelength) | 0.00410–0.5516 | | |
| Absolute scaling method | Comparison with scattering from 1 mm pure $H_2O$ | | |
| Normalization | To transmitted intensity by beam-stop counter | | |
| Monitoring for radiation damage | data frame-by-frame comparison | | |
| Exposure time | Continuous 1 s data-frame measurements of SEC elution | | |
| Sample configuration | SEC–SAXS with thermalized quartz capillary (ID 1.5mm) | | |
| Sample temperature (°C) | 20 | | |
| **Software employed for SAXS data reduction, analysis and interpretation** | | | |
| SAXS data reduction | I(q) versus q, buffer subtraction & frames selection using Foxtrot 3.10 [a] | | |
| Extinction coefficient estimate | ProtParam(Wilkins et al. 1999) | | |
| Basic analyses: Guinier, *P(r)*, MW | PRIMUSqt from ATSAS 3.2.1(Manalastas-Cantos et al. 2021) | | |
| Atomic structure modelling | Dadimodo(Rudenko et al. 2019) (https://dadimodo.synchrotron-soleil.fr/) | | |
| Missing sequence modelling | MODELLER(Webb and Sali 2014) | | |
| Three-dimensional graphic model representations | PyMOL v.0.99 | | |
| **Structural parameters** | | | |
| Guinier analysis | | | |
| I(0) ($cm^{-1}$) | 0.0184 ± 1E-4 | 0.0279 ± 1E-4 | 0.0241 ± 1E-4 |
| $R_g$ (Å) | 44.75 ± 0.31 | 48.79 ± 0.29 | 59.02 ± 0.32 |
| $q_{min}$ ($Å^{-1}$) | 0.0055 | 0.0547 | 0.00593 |
| $qR_g$ max (q min = 0.0066 $Å^{-1}$) | 1.1 | 1.29 | 1.29 |
| Coefficient of correlation, $R^2$ | 0.98 | 0.99 | 1 |
| M from Vc (ratio to predicted) | 12300 (0.85) | 168000 (1.00) | 196800 (1.1) |
| P(r) analysis | | | |
| I(0) ($cm^{-1}$) | 0.0186 ± 1E-5 | 0.0285 ± 1E-4 | 0.0248 ± 1E-4 |
| $R_g$ (Å) | 48.43 ± 0.59 | 52.88 ± 0.76 | 65.37 ± 0.68 |
| $d_{max}$ (Å) | 210 | 220 | 270 |
| q range ($Å^{-1}$) | 0.0041 to 0.50 | 0.0055 to 0.368 | 0.0059 to 0.368 |
| total estimate from GNOM | 0.67 | 0.69 | 0.67 |
| **Atomistic modelling.** | | | |
| Crystal structures | | | |
| q range for all modelling | 0.008–0.500 | 0.008–0.500 | |
| PepsiSAXS (r0 fixed) | | | |
| No constant subtraction | | | |
| $\chi^2$ | 15.62 | 3.3 | 6.05 |
| Predicted $R_g$ (Å) | 43.11 | 43.61 | 74.07 |
| Vol ($Å^3$), R0 (Å), Dro (e $Å^{-3}$) | 17944, 1.62, 0.0007 | 208111, 1.62, 0.0027 | 242115, 1.62, 0.0023 |
| Dadimodo (https://dadimodo.synchrotron-soleil.fr/) | | | |
| Starting structures | From AlphaFold2 | | |
| Rigid bodies | A: 76-170 | body1 = A: 76-172<br>body2 = A: 204-354, C: 1-408<br>body3 = A: 405-450, B: 1-226, 286-458<br>body4 = A: 476-546 | body1 = A: 76-172<br>body2 = A: 204-335, C: 1-408<br>body3 = A: 356-384, D: 60-136, E: 25-103<br>body4 = A: 405-450, B: 1-226, 286-458<br>body5 = A: 476-546 |
| No. of generated structures | 10 | 25 | 34 |
| $\chi^2$ range from PepsiSAXS | 1.49 - 2.90 | 0.65 - 1.12 | 1.09 - 1.19 |

[a] (https://www.synchrotron-soleil.fr/en/beamlines/swing#paragraphes_menu_left-block-7)

**Table S2: Yeast strains used in this study**

| Strain number | Mating type | Genotype | Reference |
|---|---|---|---|
| SL75 | h- | *ade6-704 leu1-32 ura4-D18* | Lambert et al. 2010(Lambert et al. 2010) |
| SL3456 | h+ | *pcf1-Q172A,L175A,F178A,F 179A (PIP*) ade6-704 leu1-32 ura4-D18* | This study |
| SL3452 | h+ | *pcf1-Y340A,W348A (ED*) ura4-D18 leu1-32 ade6-704* | This study |
| SL3447 | h+ | *pcf1-R147E,K150E,K154E,R161E,K168E (KER*) ura4-D18 leu1-32 ade6-704* | This study |
| SL2657 | h+ | *pcf1-477STOP (ΔWHD) ade6-704 leu1-32 ura4-D18* | This study |
| SL3233 | h+ | *FLAG:pcf1 ura4-D18 leu1-32 ade6-704* | This study |
| SL3647 | h+ | *FLAG-pcf1-PIP* ade6-704 leu1-32 ura4-D18* | This study |
| SL3650 | h+ | *FLAG-pcf1-ED* ura4-D18 leu1-32 ade6-704* | This study |
| SL3653 | h+ | *FLAG-pcf1-KER* ura4-D18 leu1-32 ade6-704* | This study |
| SL3656 | h-smt0 | *FLAG-pcf1-ΔWHD ade6-704 leu1-32 ura4-D18* | This study |
| DD122 | h+ | *pcf2:GFP:NATMX ura4-D18 leu1-32 ade6-704* | This study |
| SL3727 | h+ | *pcf2:GFP:NATMX pcf1-PIP* ura4-D18 leu1-32 ade6-704* | This study |
| SL3721 | h+ | *pcf2:GFP:NATMX pcf1-ED* ura4-D18 leu1-32 ade6-704* | This study |
| SL3724 | h+ | *pcf2:GFP:NATMX pcf1-KER* ura4-D18 leu1-32 ade6-704* | This study |
| SL3728 | h+ | *pcf2:GFP:NATMX pcf1-ΔWHD ura4-D18 leu1-32 ade6-704* | This study |
| SL3792 | h+ | *pcf2:GFP:NATMX cut11:mCherry:HYGMX ura4-D18 leu1-32 ade6-704* | This study |
| SL3602 | h+ | *rad52:GFP:KANMX pcf1::ura4+ ade6-704 leu1-32 ura4-D18* | This study |
| SL3611 | h+ | *rad52:GFP:KANMX pcf1-PIP* ade6-704 leu1-32 ura4-D18* | This study |
| SL3609 | h+ | *rad52:GFP:KANMX pcf1-ED* ade6-704 leu1-32 ura4-D18* | This study |
| SL3607 | h+ | *rad52:GFP:KANMX pcf1-KER* ade6-704 leu1-32 ura4-D18* | This study |
| SL3604 | h+ | *rad52:GFP:KANMX pcf1-ΔWHD ade6-704 leu1-32 ura4-D18* | This study |
| SL3587 | h+ | *leu1-32 ade6-704 ura4-DS/E otrR::ura4+* | This study |
| SL3589 | h+ | *pcf1::KANMX leu1-32 ade6-704 ura4-DS/E otrR::ura4+* | This study |
| SL3612 | h+ | *pcf1-PIP* leu1-32 ade6-704 ura4-DS/E otrR::ura4+* | This study |
| SL3616 | h+ | *pcf1-ED* leu1-32 ade6-704 ura4-DS/E otrR::ura4+* | This study |
| SL3624 | h+ | *pcf1-KER* leu1-32 ade6-704 ura4-DS/E otrR::ura4+* | This study |
| SL3596 | h+ | *pcf1-ΔWHD leu1-32 ade6-704 ura4-DS/E otrR::ura4+* | This study |
| VP465 | h- | *hip1::KANMX ade6-704 leu1-32 ura4-D18* | This study |

**References**


Lambert, S., K. Mizuno, J. Blaisonneau, S. Martineau, R. Chanet, K. Freon, J. M. Murray, A. M. Carr and G. Baldacci (2010). "Homologous recombination restarts blocked replication forks at the expense of genome rearrangements by template exchange." Mol Cell **39**(3): 346-359.10.1016/j.molcel.2010.07.015, PMID: 20705238

Manalastas-Cantos, K., P. V. Konarev, N. R. Hajizadeh, A. G. Kikhney, M. V. Petoukhov, D. S. Molodenskiy, A. Panjkovich, H. D. T. Mertens, A. Gruzinov, C. Borges, C. M. Jeffries, D. I. Svergun and D. Franke (2021). "ATSAS 3.0: expanded functionality and new tools for small-angle scattering data analysis." Journal of Applied Crystallography **54**: 343-355.10.1107/S1600576720013412, PMID: WOS:000613988600036

Rudenko, O., A. Thureau and J. Perez (2019). "Evolutionary refinement of the 3D structure of multi-domain protein complexes from Small Angle X-ray Scattering data." Proceedings of the 2019 Genetic and Evolutionary Computation Conference Companion (Geccco'19 Companion): 401-402.10.1145/3319619.3322002, PMID: WOS:000538328100200

Thureau, A., P. Roblin and J. Perez (2021). "BioSAXS on the SWING beamline at Synchrotron SOLEIL." Journal of Applied Crystallography **54**: 1698-1710.10.1107/S1600576721008736, PMID: WOS:000727770700016

Webb, B. and A. Sali (2014). "Protein Structure Modeling with MODELLER." Protein Structure Prediction, 3rd Edition **1137**: 1-15.10.1007/978-1-4939-0366-5_1, PMID: WOS:000333982100002

Wilkins, M. R., E. Gasteiger, A. Bairoch, J. C. Sanchez, K. L. Williams, R. D. Appel and D. F. Hochstrasser (1999). "Protein identification and analysis tools in the ExPASy server." Methods Mol Biol **112**: 531-552.10.1385/1-59259-584-7:531, PMID: 10027275
